# Characterization of glucosyltransferase and glucuronosyltransferase family members reveals how major flavone glycoside accumulate in the root of *Scutellaria baicalensis*

**DOI:** 10.1101/2021.07.24.453627

**Authors:** Tianlin Pei, Tian Li, Xiaoqiang Li, Yijia Yin, Mengying Cui, Yumin Fang, Jie Liu, Yu Kong, Ping Xu, Qing Zhao

## Abstract

Flavonoid glycosides extracted from roots of *Scutellaria baicalensis* exhibit strong pharmaceutical effect in antitumor, antioxidative, anti-inflammatory, and antiviral activity. UDP glycosyltransferase family members are responsible for the transfer of a glycosyl moiety from UDP sugars to a wide range of acceptor flavonoids. Here, we report the phylogenetic analysis, tissue-specific expression and biochemical characterization of 10 glucosyltrasferases (SbUGTs) and 6 glucuronosyltransferases (SbUGATs) based on the recently released genome of *S. baicalensis*. These results reveal that the high expression level and affinity to substrate of SbUGAT4 make baicalin become the richest flavonoid glycoside in the root of *S. baicalensis*.

---

*Scutellaria baicalensis* Georgi is an important medicinal plant used in China and other Asian countries for treatment of inflammation, diarrhea, lung and liver infections (Zhao et al., 2016a). The extracts from *S. baicalensis* were recently reported to inhibit growth of a range of cancer cells (Wang et al., 2018). Flavones in roots of *S. baicalensis* are the major bioactive compounds that responsible for these bioactivities. Around 100 flavones were reported from *S. baicalensis* roots and glycosylation contributes dramatically to the diversity of the flavone structures (Wang, et al. 2018). Glycosylation increases solubility of the flavones (compared to their aglycones), which can be easily absorbed by human body (Goel et al., 2021).

Our metabolome analysis showed that there were 69 glycosides totally found in *S. baicalensis* root tissues, with 62 *O*-glycosides and 7 *C*-glycosides (Supplementary Table 1). There were 33 flavonoid 7-*O*-glycosides (Supplementary Figure 1), which included 17 glucose moieties, 8 glucuronic acids, 3 rutinose, 2 malonylglucose, 1 galactose, 1 rhamnose, and 1 uncertain hexose, indicating that glucosylation and glucuronidation are two major glycosylated decorations in the root of *S. baicalensis*.

*O*-glycosylation in plants is catalyzed by UDP glycosyltransferase that transfer a glycosyl group to a hydroxyl group of flavones. The activity of glucuronosyltransferase (UBGAT) in *S. baicalensis* was firstly described in 2000 (Nagashima et al., 2000) and a glucosyltransferase (UBGT) from *S. baicalensis* was also characterized at molecular level that converts baicalein to oroxin A (Hirotani et al., 2000). However, in most angiosperm species, glycosyltransferases happen as gene families (Yonekura-Sakakibara and Hanada 2011). It is still unclear if there is other glucosyltransferase or glucuronosyltransferase member participates in flavone biosynthesis pathways and do they plays different roles to different compounds. With the availability of *S. baicalensis* genome, it is necessary to carry out genome-wide study of the glucosyltransferases and glucuronosyltransferases in the medicinal plant and to elucidate the relationship between the enzymes and the metabolic accumulation patterns in the plant.

We identified 10 glucosyltrasferases (*SbUGT1-10*) and 7 glucuronosyltransferase (*SbUGAT1-6*) candidate genes which might be involved in the flavonoid 7-*O* glycosylation (Supplementary Table 2 and Supplementary Figure 2). SbUGAT1.1 and SbUGAT1.2 were two distinct open reading frames predicted for a same gene locus. The full-length cDNAs of all the *SbUGTs* and *SbUGATs* were successfully isolated using specific primers (Supplementary Table 3). The genes were then reconstructed to a prokaryotic expression vector. Amino acid sequences alignment showed that the enzymes all possessed the conserved PSPG motif, and differences of two amino acids (Trp and Arg in SbUGAT) account for the functional divergence of UGT and UGAT (Supplementary Figure 3) (Noguchi et al., 2009). The SbUGTs and SbUGATs were clearly separated in phylogenetic tree (Figure 1a). SbUGT1 clustered with SbUGT2 and 3, which was the sister group of SbUGATs clade. SbUGT4, 5 and 6 comprised a subgroup, while SbUGT7, 8, 9 and 10 clustered together. Expression patterns of the *SbUGT* and *SbUGAT* genes were analyzed using FPKM values from RNA-seq data (Zhao et al., 2019). As shown in Figure 1b, *SbUGT1*, *2*, *3*, *7*, *8*, *9*, and *SbUGAT1*, *2*, *4* had relatively higher expression in both roots and MeJA-induced roots, suggesting these genes could be involved in biosynthesis of flavonoid glycosides in roots. Transcripts of *SbUGT4*, *5*, *6*, *10*, and *SbUGAT3, 5* were highly accumulated in stems, leaves and flowers, while *SbUGAT6* seemed to be a flower buds specific gene, which was probably involved in decorations of flower pigments.

**Figure 1.**
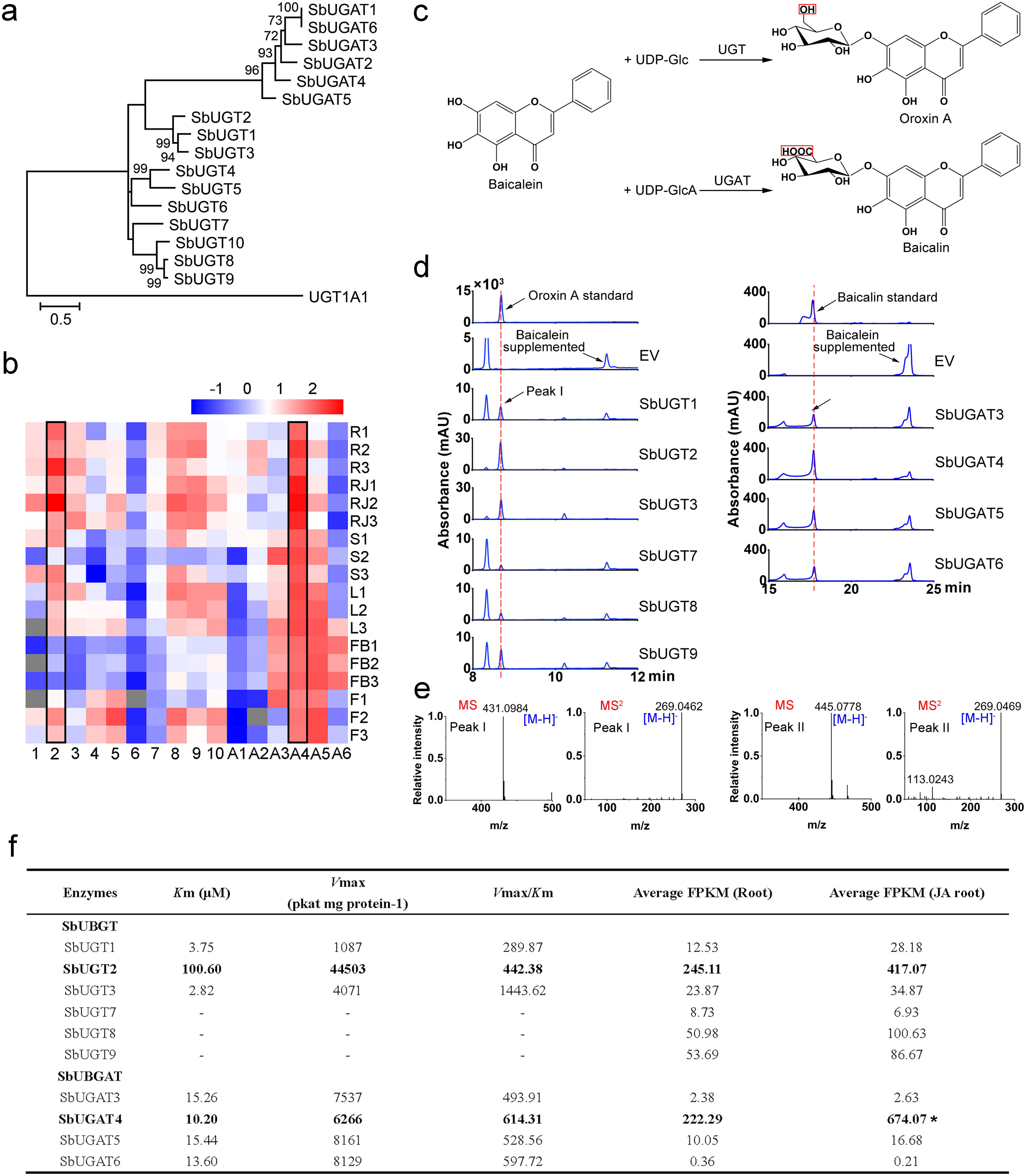
Identification and characterization of SbUGT and SbUGAT in *S. baicalensis*. a. Phylogenetic tree of SbUGT and SbUGAT proteins. Maximum-Likelihood method was used to construct the tree with bootstrap (n=1000). A glucuronosyltransferase (UGT1A1) from *Homo sapiens* was used as an outgroup. b. Tissues-specific expression heat map of the functional *SbUGT* and *SbUGAT* genes. The FPKM values of expression levels were normalized by log10, and the scale was shown top. R, root; RJ, root treated with MeJA; S, stem; L, leaf; FB, flower bud; F, flower; the numbers behind indicated replicates. Numbers on the bottom represented SbUGT1 (1), SbUGT2 (2), SbUGT3 (3), SbUGT4 (4), SbUGT5 (5), SbUGT6 (6), SbUGT7 (7), SbUGT8 (8), SbUGT9 (9), SbUGT10 (10), SbUGAT1 (A1), SbUGAT2 (A2), SbUGAT3 (A3), SbUGAT4 (A4), SbUGAT5 (A5), SbUGAT6 (A6). The columns with black boxes represented the SbUGT and SbUGAT which had the highest FPKM values of expression levels in root. c. The reaction catalyzed by SbUBGT and SbUBGAT using baicalein as a substrate. d. HPLC analysis of purified proteins of SbUGT (left column) and SbUGAT (right column) using baicalein as a substrate *in vitro*. e. MS and MS^2^ patterns of the peak I and peak II products, which were identical to oroxin A and baicalin standards, respectively. f. Kinetic parameters of SbUGT and SbUGAT incubated baicalein as a substrate, as well as their average FPKM values of root and MeJA-induced root. One asterisk (*) indicates a significant difference (0.01<*P*<0.05) between the root and MeJA-induced root under *t*-test. The products of SbUGT7, SbUGT8 and SbUGT9 were very low or not detected at linear reaction stage.

Crude proteins of these SbUGTs and SbUGATs were extracted from *E. coli* and were incubated with baicalein as substrate. As shown in Figure 1c and 1d, *in vitro* enzyme assays indicated that 6 SbUGTs (SbUGT1, SbUGT2, SbUGT3, SbUGT7, SbUGT8 and SbUGT9) could use uridine diphosphate glucose (UDP-Glc) but not UDP-glucuronic acid (UDP-GlcA) as a sugar donor and convert baicalein to oroxin A (baicalein 7-*O*-glucoside). Correspondingly, 4 SbUGATs (SbUGAT3, SbUGAT4, SbUGAT5 and SbUGAT6) used only UDP-GlcA as the sugar donor and catalyzed baicalein to baicalin (baicalein 7-*O*-glucuronide). The products were determined by comparing their retention time and MS/MS patterns with corresponding authentic standards. (Figure 1d).

The recombinant enzymes were purified from crude proteins that exhibited glycosyltransferase activity towards baicalein for kinetic analysis (Supplementary Figure 3). SbUGT3 had the lowest *K*m (2.82 μM) among all the SbUGTs, but the highest *V*max value was detected for SbUGT2, which is 44503 pkat mg^−1^ protein, leading to a 2.26-fold higher *V*max/*K*m for SbUGT3 than for SbUGT2. However, SbUGAT2 has the most abundant transcripts in root compared with other SbUGTs, with its FPKM value 10.27 times as that of SbUGT3. The lowest SbUGATs *K*m value was found for SbUGAT4, this enzyme also had the highest *V*mas/*K*m ratio, suggesting SbUGAT4 is the most efficient glucuronosyltransferase in *S. baicalensis*. Furthermore, SbUGAT4 is also the most highly expressed SbUGAT gene in root tissues. In *S.baicalensis* root, SbUGT2 and SbUGAT4 has the comparable expression level, with their FPKM 245.11 and 222.29, respectively, which is significantly higher than that of other *SbUGT* or *SbUGAT* genes. So competition between SbUGT2 and SbUGAT4 would determine the metabolic patterns in *S.baicalensis* root. As SbUGAT4 had a higher *V*max/*K*m value for baicalein, being 1.4 times higher than that of SbUGT2, this explains the large amount accumulation of baicalin, rather than oxinin A, in *S.baicalensis* root. While SbUGT2 and the other SbUGTs might be involved in biosynthesis of 4′-hydroxylated flavone 7-*O*-glucosides, such as luteolin 7-*O*-glucoside and apigenin 7-*O*-glucoside.

Specialized metabolites from plants are powerful weapons for human when challenged by a pandemic, like the COVID-19 virus which has caused 190 million people infected and over 4 million people died as we prepared this manuscript (https://covid19.who.int/). Baicalein from root of *S. baicalensis* exhibited excellent performance in suppressing replication of COVID-19 virus (Liu et al., 2020, Su et al., 2020). Baicalin is the 7-*O* glucuronidated product of baicalein converted by UGAT. The sugar moiety contributes stronger absorptivity in human intestine and the absorbed baicalein can be released from baicalin by hydrolase of human beings. Newly developed genome sequencing technologies helped us elucidated the specially-evolved pathway for baicalein (Zhao et al., 2018, Zhao, et al. 2019, Zhao et al., 2016b). For the final biosynthetic step of baicalin, 6 SbUGTs and 4 SbUGATs cloned in this study showed 7-*O*-glucosylated and 7-*O*-glucuronidated activities to baicalein. Highly expression in roots and more preferred to the substrate of SbUGAT4 make baicalin become the richest flavonoid glycoside in the root of *S. baicalensis*. Furthermore, the biosynthesis of baicalein directly from glucose *in vitro* has been achieved by *E.coli* fed-batch fermentation and the production reached to 214.1 mg/L (Ji et al., 2021). Our results provided a toolkit for the biosynthesis of baicalin by using synthetic biology.

## Authors′ contributions

T.L.P. and Q.Z initiated the program and coordinated the project. T.L.P., T.L., X.Q.L., and Y.J.Y. isolate the genes and characterized the enzymes. Y. K. assisted with the LC-MS analysis. All the authors analyzed and interpreted the data. T.L.P. wrote the manuscript. Q.Z. revised the manuscript. All authors read and approved the final manuscript.

## Acknowledgements

This work was supported by National Key R&D Program of China (2018YFC1706200), National Natural Science Foundation of China (31870282 and 31700268), Chenshan Special Fund for Shanghai Landscaping Administration Bureau Program (G182401, G192419 and G212401), and Youth Innovation Promotion Association of Chinese Academy of Sciences. QZ is also supported by the Shanghai Youth Talent Support Program and SANOFI-SIBS scholarship.

