## Supplemental Figures&Tables for "Characterization of glucosyltransferase and glucuronosyltransferase family members reveals how major flavone glycoside accumulate in the root of *Scutellaria baicalensis*"

### Materials and methods

#### Metabolome analysis

Root samples were collected from 2-month old and 2-year old *S. baicalensis* plants maintained in Shanghai Chenshan Botanical Garden, and ground into powder in liquid nitrogen then freeze dried. 20 mg of each sample was suspended in 2 ml 70% methanol and then extracted in an ultrasonic water bath for 2 h. After centrifugation at 12,000 g for 10 min, the supernatant was filtered through a 0.2 μm Millipore filter before metabolite analysis.

Samples were analyzed using an UPLC-ESI-MS/MS system comprised of UPLC, Shim-pack UFLC SHIMADZU CBM30A, (www.shimadzu.com.cn/); MS, Applied Biosystems 6500 Q TRAP, (www.appliedbiosystems.com.cn/). The analytical conditions were as follows: UPLC: column, Waters ACQUITY UPLC HSS T3 C18 (1.8 µm, 2.1 mm*100 mm), the mobile phase consisted of solvent A, pure water with 0.04% acetic acid, and solvent B, acetonitrile with 0.04% acetic acid. Sample measurements were performed with a gradient program that employed the starting conditions of 95% A, 5 % B. Within 10min, a linear gradient to 5% A, 95% B was programmed, and a composition of 5% A, 95% B was kept for 1 min. Subsequently, a composition of 95% A, 5.0 % B was applied within 0.10 min and maintained for 2.9 min. The column oven was set to 40 ℃; the injection volume was 2 μl. Alternatively, the effluent was connected to an ESI-triple quadrupole-linear ion trap (Q TRAP)-MS.

LIT and triple quadrupole (QQQ) scans were acquired on a triple quadrupole-linear ion trap mass spectrometer (Q TRAP), API 6500 Q TRAP UPLC/MS/MS System, equipped with an ESI Turbo Ion-Spray interface, operating in positive and negative ion mode and controlled by Analyst 1.6.3 software (AB Sciex). The ESI source operation parameters were as follows: ion source, turbo spray; source temperature 550 ℃; ion spray voltage (IS) 5500 V (positive ion mode)/-4500 V (negative ion mode); ion source gas I (GSI), gas II(GSII), curtain gas (CUR) were set at 50, 60, and 30.0 psi, respectively; the collision gas (CAD) was high. Instrument tuning and mass calibration were performed with 10 and 100 μmol/L polypropylene glycol solutions in QQQ and LIT modes, respectively. QQQ scans were acquired as

MRM (Multiple Reaction Monitoring) experiments with collision gas (nitrogen) set to 5 psi. DP (Declustering Potential) and CE (Collision Energy) for individual MRM transitions was done with further DP and CE optimization. A specific set of MRM transitions were monitored for each period according to the metabolites eluted within this period.

### Genome-wide identification of *SbUGT* and *SbUGAT* genes

The hidden Markov model (HMM) profile of Pfam PF00201 (http://pfam.xfam.org/) was used to extract full-length glycosyltransferase candidates from the *S. baicalensis* genome by the HMM algorithm (HMMER) (Eddy 1998), filtering by a length between 200 and 600 amino acids.

Multiple sequence alignments and phylogenetic tree construction were performed using MEGA X (Kumar et al., 2018). For the Neighbor-Joining tree, candidates were constructed under the default parameters with UGT sequences from *Arabidopsis thaliana* (downloaded from http://www.p450.kvl.dk/UGT.shtml), and with known UGTs under the following accession numbers: BpUGAT (AB190262), SlUGT1 (AB362989), AmUGTcg10 (AB362988), PfUGT50 (AB362991), SiUGT23 (AB362990), VvGT5 (AB499074), and Sb3GT1 (MK577650). 7-*O* SbUGT and SbUGAT candidates could be screened according to the annotated function and classified subfamilies. Maximum-Likelihood tree was constructed under the default parameters with sequences of 7-*O* SbUGT and SbUGAT candidates.

### Gene cloning

The complete open reading frames (ORFs) of the *SbUGT* and *SbUGAT* genes were amplified by RT-PCR using the primers listed in Supplementary Table 3. cDNA templates were chosen according to the tissue-specific expression patterns of *SbUGT* and *SbUGAT* genes. The ORFs of *SbUGT1* and *SbUGT10* were obtained by *de novo* synthesis (GenScript, Nanjing, China). According to the manufacturer’s instructions, fragments were cloned into the entry vector pDONR207 and prokaryotic expression vector pYesdest17 using the Gateway BP Clonase II Enzyme Kit and LR Clonase II Enzyme Kit (Invitrogen, MA, USA), respectively.

**Crude enzymes extraction and protein purification**

The successfully constructed vectors were transformed into *E. coli* Rosetta (DE3) competent cells (Weidi Biotech, Shanghai, China). After growing at 37 ℃ for 12 h, transformant colonies were initially grown in 10 ml of LB liquid medium with 100 mg/mL ampicillin at 37 ℃ and 180 rpm for approximately 12 h, and then transferred in 200 mL LB liquid medium with 100 mg/mL ampicillin at 37℃ in a shaking incubator until the $OD_{600}$ reached 0.6–0.8. Isopropyl β-D-thiogalactopyranoside (IPTG) was added to a final concentration of 1 mM, and cultured at 16℃ and 120 rpm for 16 h. pET28a-transformed *E. coli* Rosetta (DE3) was set as a control.

For crude enzymes extraction, *E.coli* cells were harvested by centrifugation at 12,000 rpm, and then resuspended in 50 mM phosphate buffer (pH 8.0) that contained 0.5 mM phenylmethanesulfonylfluoride (PMSF), 300 mM NaCl, 2 mM β-mercaptoethanol. High pressure cell disruption equipment (Constant Systems, Northants, UK) was used to crush the *E.coli* cells. After centrifugation at 4℃, 12,000 rpm for 20 min, approximately 10 mL of supernatant (crude protein) was collected. An equal volume of 60% glycerin was added into the supernatant for the -80℃ storage.

For protein purification, *E.coli* cells were harvested by centrifugation at 12,000 rpm, and then resuspended in 10 mL buffer A [50 mM phosphate buffer (pH 8.0), 0.5 mM phenylmethanesulfonylfluoride (PMSF), 300 mM NaCl, 2 mM β- mercaptoethanol and 10 mM imidazole]. High pressure cell disruption equipment (Constant Systems, Northants, UK) was used to crush the *E.coli* cells. After centrifugation at 4℃, 12,000 rpm for 20 min, the supernatant was mixed with 1 mL Ni–nitrilotriacetic acid (NTA) agarose (Qiagen, Germany) and stirred at 4℃ for 1 h. The mixture was packed to a column and washed three times at 4℃ with 5 mL buffer B [50 mM phosphate buffer (pH 8.0), 0.5 mM PMSF, 300 mM NaCl, 2 mM β-mercaptoethanol and 20 mM imidazole]. The protein was eluted by 1 mL buffer C [50 mM phosphate buffer (pH 8.0), 0.5 mM PMSF, 300 mM NaCl, 2 mM β-mercaptoethanol and 250 mM imidazole], and the imidazole was removed by ultrafiltration. Protein concentrations were determined using the Bradford method (Bradford 1976) and analyzed by 10% SDS-polyacrylamide gel electrophoresis

***In vitro* enzyme assays and kinetic studies**

Crude enzyme assays were performed in a 100 μl reaction volume, which contained 100 mM Tris-HCl buffer (pH 7.0), 0.5 mM sugar donor (UDP-glucose or UDP-glucuronic acid), 5 μl of extracted protein and 100 μM substrate. The reaction was incubated for 2 h at 37 ℃. Methanol was then added at a final concentration of 70% to quench the reaction. The reaction mixture was filtered with a 0.2 μm Millipore filter and analyzed by LC–MS.

For kinetics measurements, baicalein was used at concentrations ranging from 0.5 to 200 μM. Reaction time was reduced to 10 min. *K*m and *V*max values were calculated from the Eadie-Hofstee plot.

**Standard compounds**

Baicalein and baicalin were purchased from Sigma-Aldrich (St. Louis, MO, USA), and oroxin A was purchased from Yuanye-Biotech (Shanghai, China). Baicalein was dissolved in dimethyl sulfoxide (DMSO), while baicalin and oroxin A were dissolved in methanol.

**Metabolite Analyses**

Metabolites were analyzed using an Agilent 1260 Infinity II HPLC (high-performance liquid chromatography) system. Chromatographic separation was carried out on a Phenomenex Luna C18 (2) column (100×2 mm 3 μ) with a guard column. The flow rate of the mobile phase consisting of 0.1% (v/v) formic acid in water (A) and 1:1

acetonitrile/MeOH + 0.1% formic acid (B) was set to 0.26 mL/min. The gradient program was as follows: 0-3 min, 20% B; 20 min, 50% B; 20-30 min, 50% B; 36 min, 30% B; 37 min, 20% B; and 37–43 min, 20% B. The detection wavelength was 280 nm. The injection volume was 20 μl and the column temperature was 35 ℃. The products of enzyme assays were measured by comparing the area of the individual peaks with standard curves obtained from standard compounds.

LC–MS/MS was carried out by Thermo Q Exactive Plus. Chromatographic separation was carried out on a Phenomenex Luna C18 (2) column (100×2 mm 3 μ) using the same gradient described above. Mass spectra were acquired in negative ion modes with a heated ESI source, and the parameters were as follows: aus. gas flow 10 L/min; aus. Gas heater 350 ℃; sheath gas flow 40 L/min; spray voltage 3.5 kV; capillary temperature 320 ℃. For full-scan MS/data-dependent ($ddMS^2$) analysis, spectra were recorded in the m/z range of 50–750 at a resolution of 17,500 with automatic gain control (AGC) targets of $1\times10^6$ and $2\times10^5$, respectively.

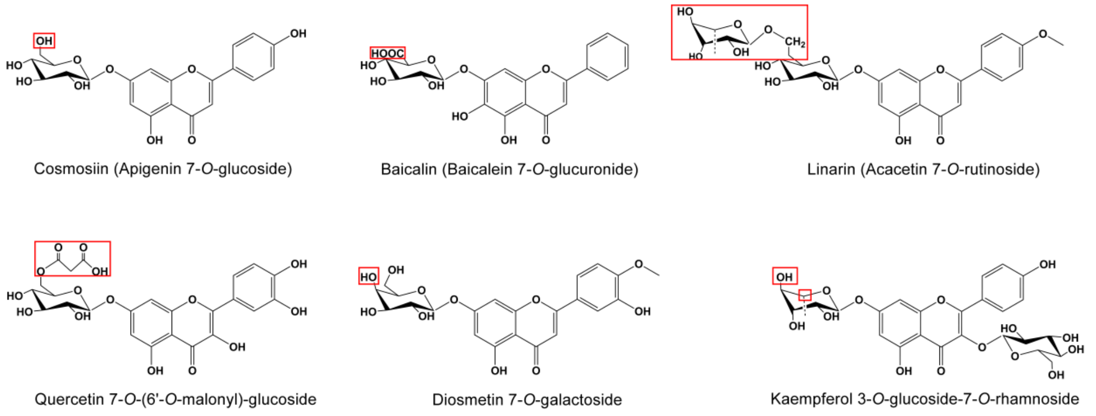


**Supplementary Figure 1. Representative 7-*O* flavonoid glycosides detected from roots of *S. baicalensis***

Red boxes indicated the different groups between sugar moieties.

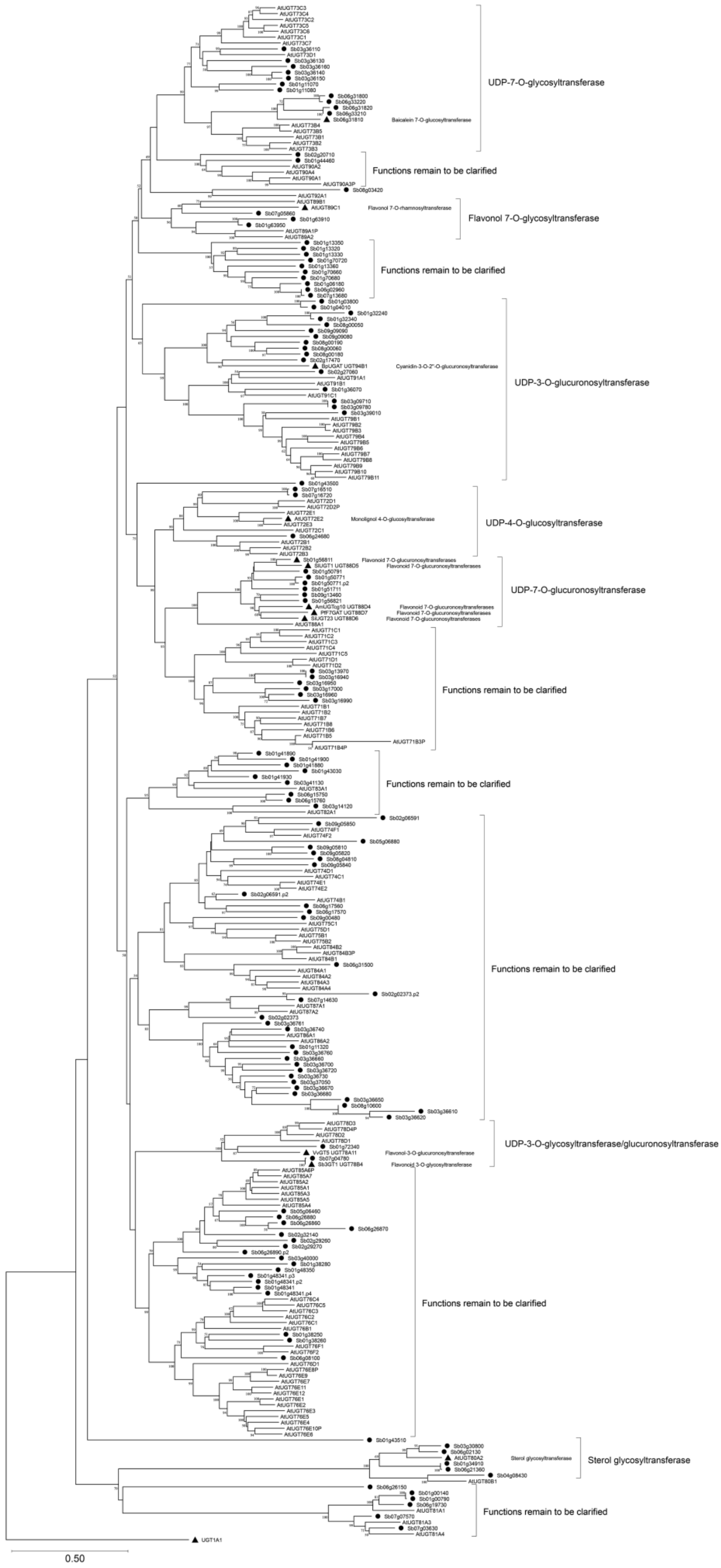


**Supplementary Figure 2. Phylogenetic tree of SbUBGT and SbUBGAT proteins**

**with UGTs from *Arabidopsis thaliana***

Neighbor-Joining method was used to construct the tree with bootstrap (n=3000). Circles before the labels represent candidate genes from *S. baicalensis*, triangles before the labels represent functional UGTs that have been reported. A glucuronosyltransferase (UGT1A1) from *Homo sapiens* was used as an outgroup.

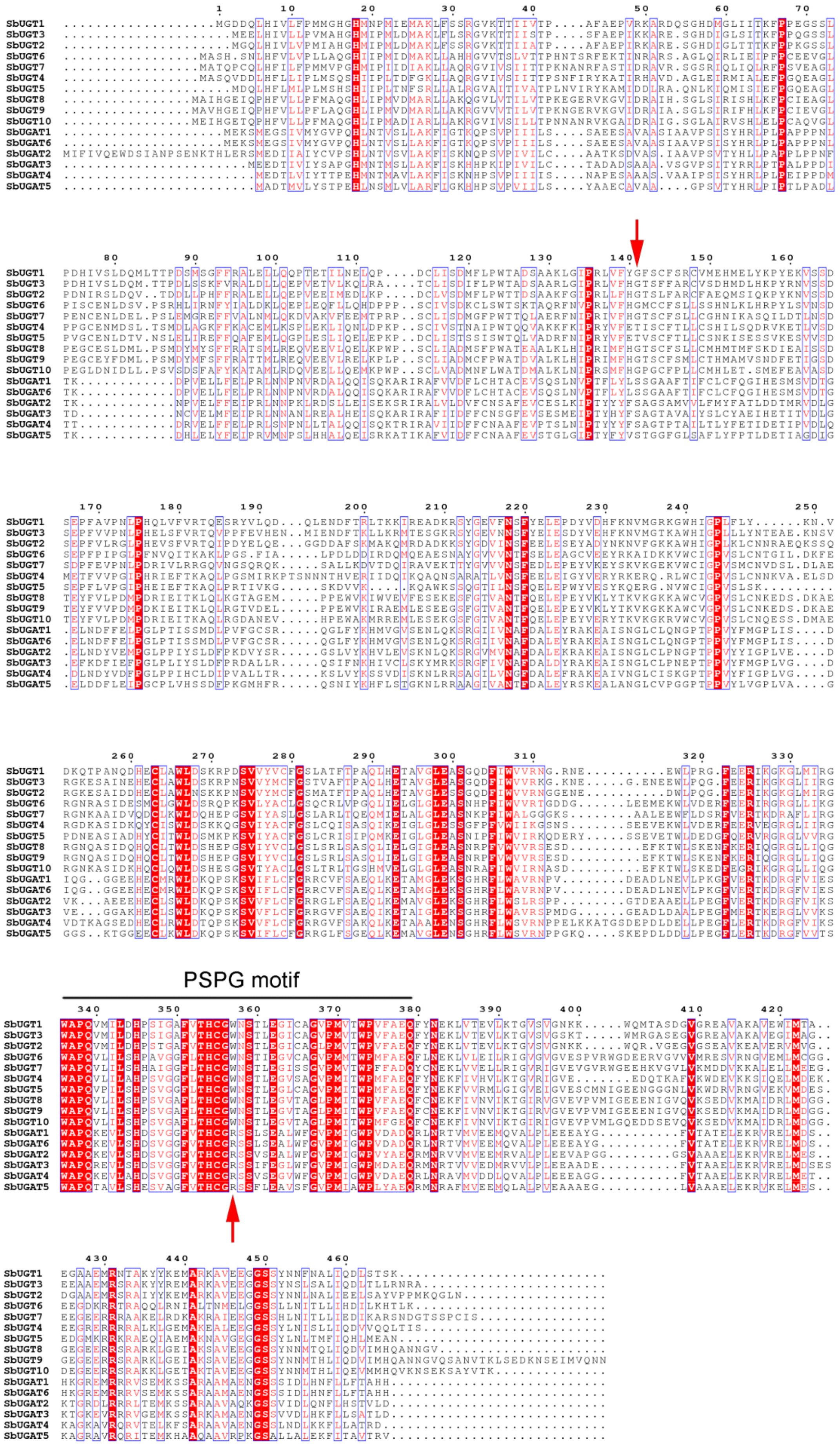
PSPG motif

**Supplementary Figure 3. Alignment of SbUGTs and SbUGATs protein sequences**
The consensus sequences were highlighted by red color. The arrows indicated the different amino acid residues between SbUGTs and SbUGATs, which were responsible for the functional divergent between these two types of glycosyltransferases.

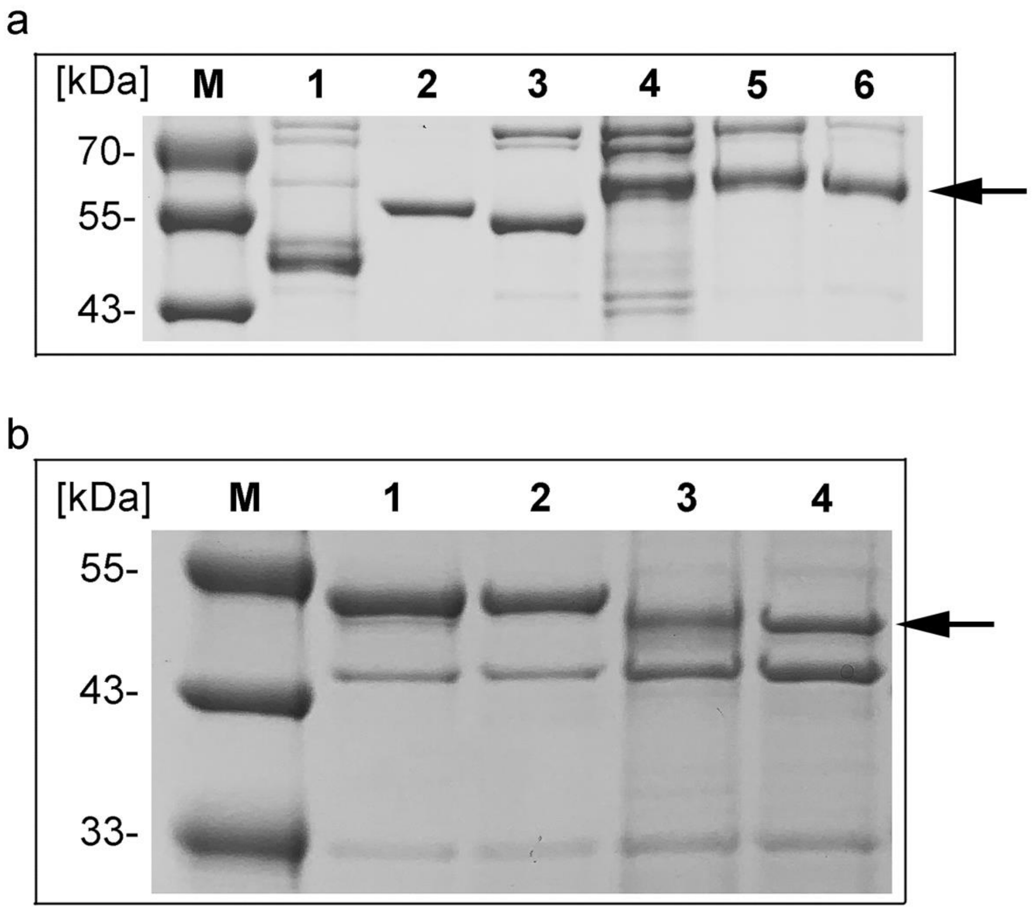


**Supplementary Figure 4. SDS PAGE analysis of purification of SbUGT and SbUGAT proteins.**

a. Tracks from left to right showed protein markers (M), SbUGT1 (1), SbUGT2 (2), SbUGT3 (3), SbUGT7 (4), SbUGT8 (5) and SbUGT9 (6).

b. Tracks from left to right showed protein markers (M), SbUGTA3 (1), SbUGAT4 (2), SbUGAT5 (3) and SbUGAT6 (4).

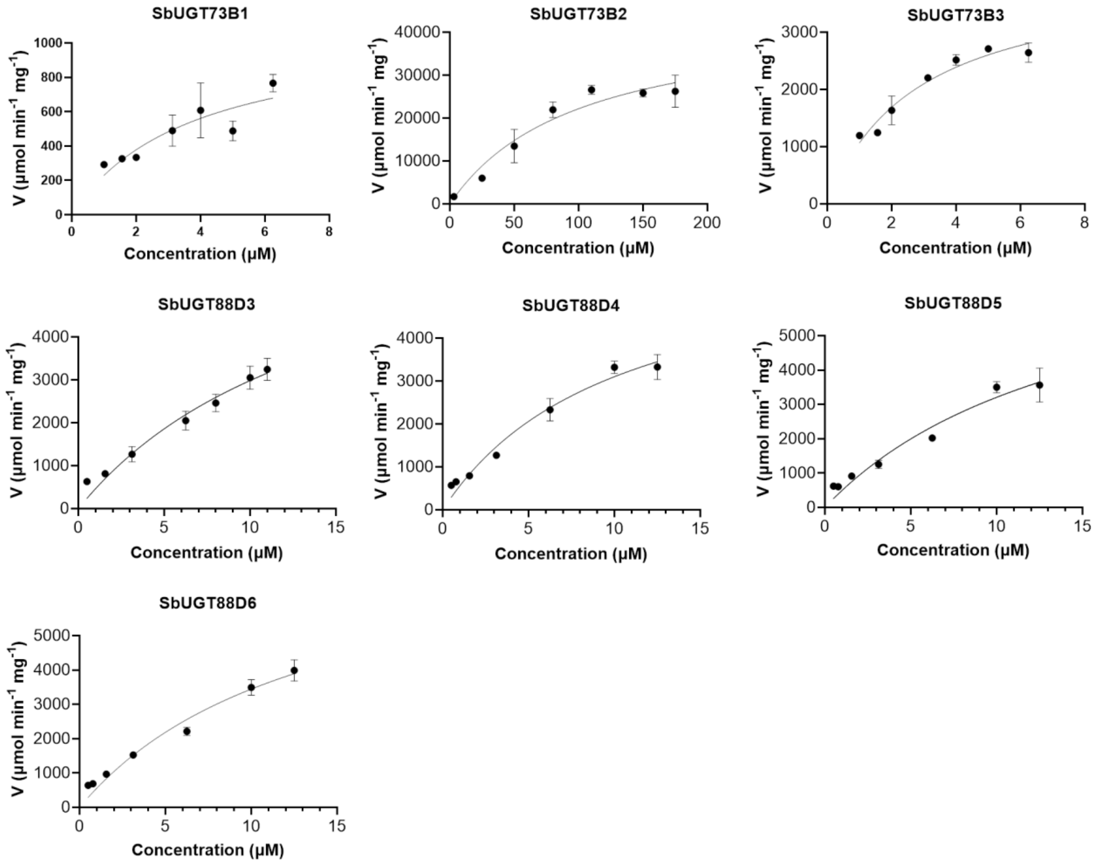


**Supplementary Figure 5. Nonlinear regressions of the Michaelis−Menten equation for SbUBGTs and SbUBGATs**

**Supplementary Table 1. Flavonoid glycosides detected in the root metabolome**

| Compounds | Molecular Weight (Da) | Formula | Ionization model | Class | CAS | R1 | R2 | R3 | YR1 | YR2 | YR3 | mix01 | mix02 | mix03 |
|---|---|---|---|---|---|---|---|---|---|---|---|---|---|---|
| Baicalin | 446.07 | C21H18O11 | [M+H]+ | Flavonoid | 21967-41-9 | 1.85E+08 | 1.80E+08 | 1.67E+08 | 2.19E+08 | 1.16E+08 | 1.76E+08 | 1.64E+08 | 1.65E+08 | 1.57E+08 |
| Apigenin 5-*O*-glucoside | 432.09 | C21H20O10 | [M+H]+ | Flavonoid | 28757-27-9 | 9.00E+00 | 9.00E+00 | 8.86E+06 | 4.48E+07 | 1.17E+08 | 9.00E+00 | 3.97E+07 | 4.49E+07 | 3.87E+07 |
| Chrysoeriol *O*-malonylhexoside | 548.09 | C25H24O14 | [M+H]+ | Flavonoid | - | 5.53E+07 | 1.91E+07 | 1.76E+07 | 3.33E+07 | 2.67E+07 | 5.26E+07 | 4.44E+07 | 3.65E+07 | 3.28E+07 |
| Diosmetin 7-*O*-galactoside | 462.10 | C22H22O11 | [M+H]+ | Flavonoid | - | 7.79E+07 | 4.48E+07 | 3.59E+07 | 9.27E+06 | 3.70E+07 | 2.23E+07 | 3.87E+07 | 3.69E+07 | 3.30E+07 |
| Wogonoside | 460.10 | C26H20O8 | [M+H]+ | Flavonoid | - | 3.25E+07 | 3.29E+07 | 1.83E+07 | 4.07E+07 | 2.06E+07 | 2.42E+07 | 2.92E+07 | 2.85E+07 | 2.70E+07 |
| Tetahydroxyflavone 7-*O*-β-D-glucuronide | 462.06 | C21H18O12 | [M-H]- | Flavonoid | - | 3.22E+06 | 4.07E+06 | 2.38E+06 | 1.16E+07 | 8.64E+06 | 1.22E+07 | 7.39E+06 | 7.83E+06 | 8.06E+06 |
| Luteolin 7-*O*-β-D-glucuronide | 448.08 | C21H20O11 | [M-H]- | Flavonoid | - | 3.21E+06 | 4.98E+06 | 6.38E+06 | 4.89E+06 | 1.03E+07 | 9.87E+06 | 7.39E+06 | 7.37E+06 | 7.57E+06 |
| Oroxylin A 7-*O*-glucuronide | 460.10 | C22H20O11 | [M+H]+ | Flavonoid | - | 1.00E+07 | 9.51E+06 | 6.79E+06 | 8.86E+06 | 4.96E+06 | 2.58E+06 | 7.76E+06 | 7.32E+06 | 6.98E+06 |
| Luteolin 7-*O*-glucoside (Cynaroside) | 448.08 | C21H20O11 | [M+H]+ | Flavonoid | 5373-11-5 | 3.00E+06 | 4.78E+06 | 6.81E+06 | 5.50E+06 | 1.64E+07 | 1.42E+07 | 6.39E+06 | 7.11E+06 | 7.41E+06 |
| Chrysin 7-*O*-glucuronide | 430.09 | C21H18O10 | [M+H]+ | Flavonoid | - | 5.65E+06 | 7.95E+06 | 5.97E+06 | 1.12E+07 | 4.59E+06 | 6.10E+06 | 6.35E+06 | 6.09E+06 | 5.17E+06 |
| Apigenin 7-*O*-glucoside (Cosmosiin) | 432.09 | C21H20O10 | [M-H]- | Flavonoid | 578-74-5 | 2.81E+05 | 1.73E+05 | 3.41E+05 | 5.44E+06 | 9.75E+06 | 1.29E+05 | 4.67E+06 | 4.91E+06 | 4.37E+06 |
| viscidulin Ⅲ 2-*O*-β-D-glucoplyranoside | 492.13 | C23H24O12 | [M+H]+ | Flavonoid | - | 4.24E+06 | 8.83E+05 | 8.94E+06 | 2.25E+06 | 5.84E+06 | 3.01E+06 | 3.97E+06 | 3.83E+06 | 3.61E+06 |
| Scutellarin | 462.06 | C21H18O12 | [M+H]+ | Flavonoid | 27740-01-8 | 1.32E+06 | 2.00E+06 | 1.41E+06 | 6.12E+06 | 5.96E+06 | 7.91E+06 | 2.71E+06 | 3.16E+06 | 3.60E+06 |
| Luteolin 7-*O*-glucuronide | 462.06 | C21H18O12 | [M+H]+ | Flavonoid | 29741-10-4 | 1.54E+06 | 2.00E+06 | 1.18E+06 | 6.06E+06 | 5.72E+06 | 7.66E+06 | 2.73E+06 | 2.97E+06 | 3.14E+06 |
| 5, 7-dihydroxy-2-methoxyflavone 7-*O*-glucuronide | 462.08 | C21H18O12 | [M+H]+ | Flavonoid | - | 1.51E+06 | 1.85E+06 | 1.23E+06 | 6.01E+06 | 5.66E+06 | 7.28E+06 | 2.75E+06 | 3.10E+06 | 2.98E+06 |

| Compounds | Molecular Weight (Da) | Formula | Ionization model | Class | CAS | R1 | R2 | R3 | YR1 | YR2 | YR3 | mix01 | mix02 | mix03 |
|---|---|---|---|---|---|---|---|---|---|---|---|---|---|---|
| 5, 7-dihydroxy-6, 8-dimethoxyflavone 7-*O*-glucoside | 490.11 | C23H22O12 | [M+H]+ | Flavonoid | - | 5.56E+06 | 2.63E+06 | 1.98E+06 | 3.45E+06 | 1.89E+06 | 2.44E+06 | 2.98E+06 | 2.75E+06 | 2.56E+06 |
| Chrysoeriol 7-*O*-[β-D-glucuronopyranosyl-(1→2)-*O*-β-D-glucuronopyranoside] | 652.10 | C28H28O18 | [M+H]+ | Flavonoid | - | 3.95E+06 | 2.84E+06 | 1.36E+06 | 2.78E+06 | 1.29E+06 | 3.22E+06 | 2.65E+06 | 2.39E+06 | 2.32E+06 |
| Rhoifolin | 578.14 | C27H30O14 | [M+H]+ | Flavonoid | 17306-46-6 | 4.38E+06 | 2.30E+06 | 2.94E+06 | 1.27E+06 | 2.30E+06 | 1.37E+06 | 2.92E+06 | 2.31E+06 | 1.94E+06 |
| Diosmetin 7-*O*-glucuronide | 476.08 | C22H20O12 | [M+H]+ | Flavonoid | 35110-20-4 | 3.32E+06 | 3.38E+06 | 1.51E+06 | 1.72E+06 | 1.28E+06 | 1.52E+06 | 1.71E+06 | 1.73E+06 | 1.60E+06 |
| Apigenin 7-*O*-(6'-*O*-acetyl)-β-D-glucoside | 474.10 | C23H22O11 | [M+H]+ | Flavonoid | - | 1.63E+06 | 1.10E+06 | 7.50E+05 | 1.63E+06 | 7.53E+05 | 2.16E+06 | 1.48E+06 | 1.40E+06 | 1.21E+06 |
| Diosmetin 7-*O*-(6'-*O*-malonyl)-β-D-glucoside | 548.09 | C25H24O14 | [M+H]+ | Flavonoid | - | 1.31E+06 | 1.94E+06 | 2.67E+06 | 3.81E+04 | 1.05E+05 | 2.63E+05 | 1.19E+06 | 1.02E+06 | 8.78E+05 |
| Acacetin *O*-glucuronic acid | 460.08 | C22H20O11 | [M-H]- | Flavonoid | - | 5.01E+05 | 5.77E+05 | 3.68E+05 | 1.18E+06 | 1.04E+06 | 1.14E+06 | 9.08E+05 | 9.83E+05 | 9.11E+05 |
| Acacetin 7-*O*-rutinoside | 592.15 | C28H32O14 | [M+H]+ | Flavonoid | 480-36-4 | 2.82E+05 | 3.88E+05 | 1.87E+05 | 7.08E+05 | 4.33E+05 | 9.41E+05 | 5.89E+05 | 6.42E+05 | 5.55E+05 |
| Scuteamoenoside | 464.13 | C22H24O11 | [M+H]+ | Flavonoid | 123914-35-2 | 6.01E+05 | 7.58E+05 | 3.02E+05 | 8.41E+05 | 3.36E+05 | 4.02E+05 | 6.43E+05 | 5.38E+05 | 4.79E+05 |
| Chrysoeriol 5-*O*-hexoside | 462.10 | C22H22O11 | [M-H]- | Flavonoid | - | 6.70E+05 | 4.59E+05 | 6.26E+05 | 4.67E+05 | 3.80E+05 | 4.94E+05 | 4.05E+05 | 4.78E+05 | 3.64E+05 |
| Luteolin *O*-sinapoylhexoside | 654.13 | C32H30O15 | [M+H]+ | Flavonoid | - | 1.60E+05 | 9.47E+04 | 2.08E+05 | 5.25E+04 | 1.17E+06 | 4.10E+05 | 4.18E+05 | 4.55E+05 | 3.02E+05 |
| Chrysoeriol *O*-hexosyl-*O*-hexoside | 624.14 | C28H32O16 | [M+H]+ | Flavonoid | - | 1.19E+06 | 1.92E+05 | 3.76E+05 | 1.15E+05 | 1.39E+05 | 1.41E+05 | 4.36E+05 | 3.22E+05 | 3.16E+05 |
| Chrysoeriol *O*-glucuronic acid-*O*-hexoside | 638.12 | C28H30O17 | [M+H]+ | Flavonoid | - | 1.38E+06 | 2.53E+05 | 1.43E+05 | 8.79E+04 | 1.08E+05 | 1.69E+05 | 3.13E+05 | 3.13E+05 | 2.79E+05 |
| Luteolin 7, 3'-*O*-β-D-diglucoside | 610.13 | C27H30O16 | [M+H]+ | Flavonoid | 257-72-4-7 | 7.19E+04 | 1.12E+05 | 3.39E+05 | 2.95E+05 | 3.25E+05 | 4.74E+05 | 2.93E+05 | 3.13E+05 | 2.36E+05 |
| Chrysoeriol 7-*O*-hexoside | 462.10 | C22H2 | [M-H]- | Flavonoid | - | 9.97 | 2.88 | 2.46 | 9.00 | 5.78 | 1.49 | 2.44 | 2.66 | 3.03 |

| Compounds | Molecular Weight (Da) | Formula | Ionization model | Class | CAS | R1 | R2 | R3 | YR1 | YR2 | YR3 | mix01 | mix02 | mix03 |
|---|---|---|---|---|---|---|---|---|---|---|---|---|---|---|
| | | 2O11 | | | | E+05 | E+05 | E+05 | E+00 | E+05 | E+05 | E+05 | E+05 | E+05 |
| Luteolin 7-*O*-β-D-rutinoside | 594.13 | C27H30O15 | [M-H]- | Flavonoid | - | 7.16E+04 | 3.92E+04 | 8.68E+04 | 9.85E+04 | 2.32E+05 | 1.31E+05 | 1.18E+05 | 1.12E+05 | 1.09E+05 |
| Luteolin 3', 7-*O*-diglucoside | 610.13 | C27H30O16 | [M+H]+ | Flavonoid | 52187-80-1 | 3.72E+04 | 3.22E+04 | 1.06E+05 | 1.43E+05 | 1.82E+05 | 2.10E+05 | 1.03E+05 | 1.17E+05 | 1.13E+05 |
| Chrysoeriol 7-*O*-rutinoside | 608.15 | C28H32O15 | [M-H]- | Flavonoid | - | 2.18E+04 | 3.62E+04 | 1.08E+04 | 5.57E+04 | 3.92E+04 | 2.83E+04 | 3.63E+04 | 4.38E+04 | 3.47E+04 |
| Kaempferol 7-*O*-glucosdie | 448.08 | C21H20O11 | [M-H]- | Flavonols | 16290-07-6 | 1.65E+06 | 1.21E+06 | 3.94E+06 | 3.21E+06 | 2.86E+06 | 3.10E+06 | 3.08E+06 | 3.58E+06 | 2.99E+06 |
| Isorhamnetin 3-*O*-rutinoside | 624.14 | C28H32O16 | [M-H]- | Flavonols | 604-80-8 | 2.02E+05 | 3.13E+05 | 3.47E+05 | 8.98E+05 | 1.43E+06 | 1.18E+06 | 1.09E+06 | 1.05E+06 | 9.98E+05 |
| Quercetin 3, 7-*O*-β-D-diglucoside | 626.12 | C27H30O17 | [M+H]+ | Flavonols | - | 2.28E+05 | 4.96E+05 | 7.13E+05 | 4.66E+05 | 1.82E+06 | 1.40E+06 | 9.09E+05 | 7.92E+05 | 8.64E+05 |
| Syringetin 3-*O*-hexoside | 508.10 | C23H24O13 | [M+H]+ | Flavonols | - | 3.43E+05 | 2.32E+05 | 6.13E+05 | 4.76E+05 | 3.13E+06 | 5.82E+05 | 6.36E+05 | 7.73E+05 | 7.82E+05 |
| 6-hydroxykaempferol 6, 7-*O*-diglucoside | 626.12 | C27H30O17 | [M+H]+ | Flavonols | - | 1.75E+05 | 3.37E+05 | 5.97E+05 | 2.84E+05 | 7.42E+05 | 6.32E+05 | 4.99E+05 | 4.59E+05 | 4.11E+05 |
| Kaempferol 3-O-rutinoside(Nicotiflorin) | 594.13 | C27H30O15 | [M-H]- | Flavonols | 17650-84-9 | 5.75E+04 | 6.99E+04 | 8.94E+04 | 6.43E+04 | 1.75E+05 | 1.38E+05 | 1.24E+05 | 1.22E+05 | 1.26E+05 |
| Kaempferol 3-O-robinobioside(Biorobin) | 594.13 | C27H30O15 | [M-H]- | Flavonols | 17297-56-2 | 4.44E+04 | 6.53E+04 | 9.75E+04 | 9.09E+04 | 1.66E+05 | 1.36E+05 | 1.05E+05 | 1.26E+05 | 1.30E+05 |
| Quercetin 7-*O*-(6'-*O*-malonyl)-β-D-glucoside | 550.07 | C24H22O15 | [M+H]+ | Flavonols | - | 2.15E+05 | 1.13E+05 | 5.77E+04 | 7.38E+04 | 1.28E+05 | 7.40E+04 | 1.32E+05 | 1.21E+05 | 8.31E+04 |
| Kaempferol 3-*O*-glucoside-7-*O*-rhamnoside | 594.13 | C27H30O15 | [M+H]+ | Flavonols | - | 3.97E+04 | 1.01E+05 | 5.02E+04 | 4.53E+04 | 1.67E+05 | 7.78E+04 | 8.09E+04 | 7.71E+04 | 7.19E+04 |
| 6-hydroxykaempferol 3,6-*O*-diglucoside | 626.12 | C27H30O17 | [M+H]+ | Flavonols | - | 1.18E+05 | 6.36E+04 | 7.73E+04 | 1.39E+04 | 1.10E+05 | 7.83E+04 | 6.71E+04 | 5.93E+04 | 5.57E+04 |
| Isorhamnetin 3-*O*-β-(2"-*O*-acetyl-β-D-glucuronide) | 534.08 | C24H22O14 | [M-H]- | Flavonols | - | 1.62E+04 | 2.49E+04 | 2.90E+04 | 7.61E+03 | 3.92E+04 | 1.52E+04 | 4.11E+04 | 3.51E+04 | 2.51E+04 |
| Hyperin | 464.08 | C21H20O12 | [M-H]- | Flavonols | 482-36-0 | 2.40E+04 | 3.29E+04 | 3.11E+04 | 1.69E+04 | 5.53E+04 | 2.23E+04 | 2.78E+04 | 3.48E+04 | 2.21E+04 |

| Compounds | Molecular Weight (Da) | Formula | Ionization model | Class | CAS | R1 | R2 | R3 | YR1 | YR2 | YR3 | mix01 | mix02 | mix03 |
|---|---|---|---|---|---|---|---|---|---|---|---|---|---|---|
| Isorhamnetin *O*-acetyl-hexoside | 520.10 | C24H24O13 | [M-H]- | Flavonols | - | 2.12E+04 | 2.40E+04 | 2.48E+04 | 1.38E+04 | 3.83E+04 | 1.20E+04 | 2.55E+04 | 2.86E+04 | 2.46E+04 |
| Kaempferide 3-*O*-β-D-glucuronide | 476.08 | C22H20O12 | [M-H]- | Flavonols | - | 3.08E+04 | 2.93E+04 | 2.67E+04 | 4.44E+03 | 2.57E+04 | 2.01E+03 | 1.68E+04 | 1.44E+04 | 1.64E+04 |
| Eriodictyol 7-*O*-glucoside | 450.10 | C21H22O11 | [M+H]+ | Dihydroflavone | 38965-51-4 | 4.68E+06 | 7.24E+06 | 1.04E+07 | 8.71E+06 | 2.33E+07 | 1.91E+07 | 1.16E+07 | 1.13E+07 | 1.16E+07 |
| Naringenin 7-*O*-glucoside | 434.10 | C21H22O10 | [M-H]- | Dihydroflavone | 529-55-5 | 5.82E+04 | 3.79E+04 | 8.41E+04 | 1.48E+05 | 1.66E+05 | 9.11E+04 | 1.17E+05 | 1.47E+05 | 1.06E+05 |
| Naringin | 580.15 | C27H32O14 | [M-H]- | Dihydroflavone | 10236-47-2 | 9.83E+04 | 3.38E+04 | 1.24E+05 | 2.74E+04 | 2.93E+04 | 9.00E+00 | 5.63E+04 | 5.25E+04 | 8.70E+04 |
| Iridin | 522.11 | C24H26O13 | [M+H]+ | Isoflavones | 491-74-7 | 1.01E+07 | 6.51E+06 | 1.61E+07 | 3.08E+07 | 4.76E+07 | 3.90E+07 | 2.84E+07 | 2.61E+07 | 2.27E+07 |
| Calycosin 7-*O*-glucoside | 446.10 | C22H22O10 | [M+H]+ | Isoflavones | 20633-67-4 | 5.94E+06 | 1.62E+07 | 1.12E+07 | 1.26E+07 | 2.08E+07 | 2.35E+07 | 1.46E+07 | 1.41E+07 | 1.36E+07 |
| Iristectorin B | 492.11 | C23H24O12 | [M+H]+ | Isoflavones | 94396-09-5 | 1.15E+07 | 2.55E+06 | 2.57E+07 | 5.93E+06 | 1.51E+07 | 7.57E+06 | 1.30E+07 | 1.06E+07 | 9.57E+06 |
| Engeletin | 434.10 | C21H22O10 | [M-H]- | Dihydroflavonol | 572-31-6 | 2.38E+05 | 2.49E+05 | 2.50E+05 | 4.37E+05 | 2.84E+05 | 2.50E+05 | 3.84E+05 | 3.92E+05 | 3.02E+05 |
| Malvidin 3-*O*-galactoside | 493.11 | C23H25O12 | [M]+ | Anthocyanins | 30113-37-2 | 1.96E+05 | 6.76E+05 | 1.13E+06 | 9.00E+00 | 4.98E+07 | 5.72E+07 | 1.28E+07 | 1.44E+07 | 1.38E+07 |
| Malvidin 3-*O*-glucoside (Oenin) | 493.11 | C23H25O12 | [M]+ | Anthocyanins | 18470-06-9 | 1.11E+05 | 2.97E+05 | 6.17E+05 | 9.00E+00 | 2.58E+07 | 2.79E+07 | 7.09E+06 | 7.08E+06 | 7.28E+06 |
| Cyanidin 3-*O*-malonylhexoside | 534.08 | C24H22O14 | [M+H]+ | Anthocyanins | - | 1.34E+06 | 1.86E+06 | 9.00E+00 | 4.15E+06 | 3.38E+05 | 5.35E+05 | 9.76E+05 | 8.68E+05 | 1.02E+06 |
| Cyanidin 3-*O*-glucoside (Kuromanin) | 449.09 | C21H21O11 | [M]+ | Anthocyanins | 7084-24-4 | 5.02E+05 | 4.76E+05 | 9.00E+00 | 3.62E+06 | 9.00E+00 | 9.00E+00 | 7.47E+05 | 8.12E+05 | 8.16E+05 |
| Malvidin *O*-hexoside | 492.11 | C23H24O12 | [M+H]+ | Anthocyanins | - | 1.37E+04 | 3.86E+04 | 5.99E+04 | 9.00E+00 | 2.50E+06 | 2.98E+06 | 7.10E+05 | 7.59E+05 | 7.07E+05 |
| Cyanin chloride | 611.13 | C27H31O16 | [M]+ | Anthocyanins | 2611-67-8 | 6.57E+04 | 1.35E+05 | 4.33E+05 | 3.04E+05 | 1.50E+06 | 1.28E+06 | 5.58E+05 | 5.33E+05 | 5.43E+05 |
| Cyanidin 3-*O*-galactoside | 448.08 | C21H2 | [M+H]+ | Anthocyani | 27661- | 1.06 | 1.25 | 9.11 | 3.89 | 9.00 | 9.00 | 3.05 | 2.98 | 2.39 |

| Compounds | Molecular Weight (Da) | Formula | Ionization model | Class | CAS | R1 | R2 | R3 | YR1 | YR2 | YR3 | mix01 | mix02 | mix03 |
|---|---|---|---|---|---|---|---|---|---|---|---|---|---|---|
| | | 0O11 | | ns | 36-5 | E+05 | E+05 | E+05 | E+05 | E+00 | E+00 | E+05 | E+05 | E+05 |
| Malvidin 3, 5-diglucoside (Malvin) | 655.16 | $C_{29}H_{35}O_{17}$ | [M]+ | Anthocyanins | 16727-30-3 | 9.00E+00 | 9.00E+00 | 9.00E+00 | 9.00E+00 | 2.04E+05 | 2.44E+05 | 6.77E+04 | 6.77E+04 | 7.32E+04 |
| Isoschaftoside | 564.12 | $C_{26}H_{28}O_{14}$ | [M+H]+ | Flavonoid carbonoside | 52012-29-0 | 1.12E+07 | 7.15E+06 | 9.52E+06 | 1.34E+07 | 3.48E+07 | 1.88E+07 | 1.66E+07 | 1.50E+07 | 1.44E+07 |
| Apigenin 6,8-*C*-diglucoside | 594.13 | $C_{27}H_{30}O_{15}$ | [M+H]+ | Flavonoid carbonoside | 23666-13-9 | 5.73E+05 | 3.35E+05 | 7.33E+05 | 9.03E+05 | 1.94E+06 | 1.07E+06 | 9.78E+05 | 8.52E+05 | 9.05E+05 |
| Apigenin *C*-hexosyl- *O*-pentoside | 564.12 | $C_{26}H_{28}O_{14}$ | [M+H]+ | Flavonoid carbonoside | - | 4.15E+05 | 3.05E+05 | 4.45E+05 | 5.67E+05 | 1.26E+06 | 7.16E+05 | 6.06E+05 | 6.60E+05 | 5.41E+05 |
| 8-*C*-hexosyl-hesperetin-*O*-hexoside | 626.16 | $C_{28}H_{34}O_{16}$ | [M+H]+ | Flavonoid carbonoside | - | 1.26E+05 | 2.14E+05 | 3.47E+05 | 1.83E+05 | 8.99E+05 | 8.04E+05 | 4.18E+05 | 4.13E+05 | 3.89E+05 |
| 8-*C*-hexosyl-luteolin-*O*-pentoside | 580.12 | $C_{26}H_{28}O_{15}$ | [M+H]+ | Flavonoid carbonoside | - | 3.95E+05 | 2.47E+05 | 1.21E+06 | 1.05E+05 | 2.35E+05 | 1.50E+05 | 3.58E+05 | 3.44E+05 | 3.27E+05 |
| 8-*C*-hexosyl-apigenin -*O*-feruloylhexoside | 770.17 | $C_{37}H_{38}O_{18}$ | [M+H]+ | Flavonoid carbonoside | - | 7.25E+03 | 1.94E+04 | 1.68E+04 | 4.18E+04 | 4.19E+04 | 7.87E+04 | 4.05E+04 | 2.52E+04 | 2.39E+04 |
| Schaftoside | 564.12 | $C_{26}H_{28}O_{14}$ | [M+H]+ | Flavonoid carbonoside | 51938-32-0 | 2.34E+04 | 1.50E+04 | 2.24E+04 | 1.93E+04 | 8.14E+04 | 3.21E+04 | 1.94E+04 | 2.39E+04 | 1.91E+04 |

**Supplementary Table 2. The list of enzyme names, gene locus, and their subfamilies of predicted 7-*O* glycosyltransferases in *S. baicalensis***

| Enzyme names | Gene locus | Subfamily |
|---|---|---|
| **SbUBGT** | | |
| SbUGT1 | Sb06g31800 | UGT73B |
| SbUGT2 | Sb06g31810 | UGT73B |
| SbUGT3 | Sb06g31820 | UGT73B |
| SbUGT4 | Sb01g11070 | UGT73C |
| SbUGT5 | Sb01g11080 | UGT73C |
| SbUGT6 | Sb03g36110 | UGT73D |
| SbUGT7 | Sb03g36130 | UGT73D |
| SbUGT8 | Sb03g36140 | UGT73D |
| SbUGT9 | Sb03g36150 | UGT73D |
| SbUGT10 | Sb03g36160 | UGT73D |
| **SbUBGAT** | | |
| SbUGAT1.1 | Sb01g50771.p1 | UGT88D |
| SbUGAT1.2 | Sb01g50771.p2 | UGT88D |
| SbUGAT2 | Sb01g50791 | UGT88D |
| SbUGAT3 | Sb01g51711 | UGT88D |
| SbUGAT4 | Sb01g56811 | UGT88D |
| SbUGAT5 | Sb01g56821 | UGT88D |
| SbUGAT6 | Sb09g13460 | UGT88D |

**Supplementary Table 3. Primers used for the cloning of *SbUGT* and *SbUGAT* genes**

| Gene names | Forward (5′-3′) | Reverse (5′-3′) |
|---|---|---|
| SbUGT2 | GGGGACAAGTTTGTACAAAAAAGCAGGCTTCATGGGACAACTCCACATAGTCC | GGGGACCACTTTGTACAAGAAAGCTGGGTTTTAGTTTAAGCCCTGTTTCATAGG |
| SbUGT3 | GGGGACAAGTTTGTACAAAAAAGCAGGCTTCATGGAAGAGCTACATATTGTCCTTC | GGGGACCACTTTGTACAAGAAAGCTGGGTTTTATGCCCTGTTTCTTAGGAGTG |
| SbUGT4 | GGGGACAAGTTTGTACAAAAAAGCAGGCTTCATGGCTTCCCAAGTTGATGA | GGGGACCACTTTGTACAAGAAAGCTGGGTTTCAAGAAATAGTAAGTTGTTGCAC |
| SbUGT5 | GGGGACAAGTTTGTACAAAAAAGCAGGCTTCATGGATCAACTCCACTTTCTTATG | GGGGACCACTTTGTACAAGAAAGCTGGGTTTTAATTTGCTTCCATGAGATGTTG |
| SbUGT6 | GGGGACAAGTTTGTACAAAAAAGCAGGCTTCATGGCGTCCCATTCAAACCT | GGGGACCACTTTGTACAAGAAAGCTGGGTTCTATTTCAAGGTGTGTTTCAAAA |
| SbUGT7 | GGGGACAAGTTTGTACAAAAAAGCAGGCTTCATGGCCACTCAACCTTGCC | GGGGACCACTTTGTACAAGAAAGCTGGGTTTCAAGAAATGCAAGGTGATGAT |
| SbUGT8 | GGGGACAAGTTTGTACAAAAAAGCAGGCTTCATGGCAATCCATGGAGAAATAC | GGGGACCACTTTGTACAAGAAAGCTGGGTTTTAGACCCCATTATTGGCCT |
| SbUGT9 | GGGGACAAGTTTGTACAAAAAAGCAGGCTTCATGGCAGTCCATGGAGAAATAC | GGGGACCACTTTGTACAAGAAAGCTGGGTTTCAATTATTCTGAACCATAATTTCA |
| SbUGAT1 | GGGGACAAGTTTGTACAAAAAAGCAGGCTTCATGGAAAAATCAATGGAAGGCA | GGGGACCACTTTGTACAAGAAAGCTGGGTTTCAATGGTGGGCAGTAAAGAGA |
| SbUGAT2 | GGGGACAAGTTTGTACAAAAAAGCAGGCTTCATGATTTTTATAGTGCAGGAATGGG | GGGGACCACTTTGTACAAGAAAGCTGGGTTTTAATCAAGGACGGTGGAGTGAA |
| SbUGAT3 | GGGGACAAGTTTGTACAAAAAAGCAGGCTTCATGGAAGAAGACACCATTGTAATT | GGGGACCACTTTGTACAAGAAAGCTGGGTTTCAATCAAGGGTGGCGGAGA |
| SbUGAT4 | GGGGACAAGTTTGTACAAAAAAGCAGGCTTCATGGAAGACACACTTGTGATCTACA | GGGGACCACTTTGTACAAGAAAGCTGGGTTTTAATCCCGAGTGGCGAGAAG |
| SbUGAT5 | GGGGACAAGTTTGTACAAAAAAGCAGGCTTCATGGCGGACACCATGGTTC | GGGGACCACTTTGTACAAGAAAGCTGGGTTTTAAACCCGAGTCACCGCC |

| Gene names | Forward (5′-3′) | Reverse (5′-3′) |
|---|---|---|
| SbUGAT6 | GGGGACAAGTTTGTACAAAAAAGCAGGCTTCATGGAAGACACCATTGTTCTGTATG | GGGGACCACTTTGTACAAGAAAGCTGGGTTTTAATCCCGAGTGGCGAGAAG |
